# LiPLike: Towards gene regulatory network predictions of high-certainty

**DOI:** 10.1101/651596

**Authors:** Rasmus Magnusson, Mika Gustafsson

## Abstract

**Motivation:** Reverse engineering of gene regulatory networks has for years struggled with high correlation in expression between regulatory elements. If two regulators have matching expression patterns it is impossible to differentiate between the two, and thus false positive identifications are abundant.

**Results:** To allow for gene regulation predictions of high confidence, we propose a novel method, LiPLike, that assumes a regression model and iteratively searches for interactions that cannot be replaced by a linear combination of other predictors. To compare the performance of LiPLike with other available inference methods, we benchmarked LiPLike using three independent datasets from the previous DREAM5 challenge. We found that LiPLike could be used to stratify predictions of other inference tools, and when applied to the predictions of DREAM5 participants we observed the accuracy to on average be improved *>*140% compared to individual methods. Furthermore, we observed that LiPLike independently predicted networks better than all DREAM5 participants when applied to biological data. When predicting the *Escherichia coli* network, LiPLike had an accuracy of 0.38 for the top-ranked 100 interactions, whereas the corresponding DREAM5 consensus model yielded an accuracy of 0.11.

**Availability:** We made LiPLike available to the community as a Python toolbox, available at https://gitlab.com/Gustafsson-lab/liplike. We believe that LiPLike will be used for high confidence predictions in studies where individual model interactions are of high importance, and that LiPLike will be used to remove false positive predictions made by other state-of-the-art gene-gene regulation prediction tools.

## 1 Introduction

Understanding and interpreting big data has become a focal point of the field of bioinformatics, fueled by an increasing stream of data from several omics’ techniques. The size and complexity of data from modern state-of-the-art techniques such as RNA-Seq expression at first overwhelmed researchers, as few conclusions can be drawn from simple investigations [1]. Nevertheless, network analysis has emerged as a prominent tool that is used to both distinguish and interpret cellular processes [2–6]. This network analysis can be performed in several ways, such as by constructing graphs of nodes and edges that give information about cellular processes. When reverse engineering gene regulatory networks (GRNs), genes are denoted as nodes while the aim is to infer interactions between them, referred to as edges. Next, studying the GRN can give different insights into the biological system [7]. These insights have been shown to include identifications of important feed-back or cross-talks in biological systems [1], understanding of upstream master regulators of gene expression [8, 9], and potential drug targets [7, 10].

Currently, there are several methods used to predict GRNs from gene expression data. A common approach is to assume the data to be generated from a linear system and apply regression-based algorithms. Such methods include the LASSO [11], the Elastic Net [12], and the Inferelator [13, 14]. Other popular methods include mutual information, notably ARACNe [15], Bayesian modelling [16, 17], neural networks [18], and methods based on correlation [19], all of which are methods that aim at reconstructing GRNs.

Although there are a number of cases where GRN inference has been successfully applied to solve biological questions [7, 8, 10, 20], network analysis has historically struggled with a set of problems. First, the results of network inference are often dependent on the underlying quality of data, with factors such as correlated regulators posing a problem for most algorithms [21–23]. Second, understanding what aspects of the results can be trusted often poses a problem, as most GRN inference methods aim to recreate the most probable network, which often comes at the expense of a high false prediction rate. An example of these low accuracies can be found in the prestigious DREAM5 GRN prediction challenge, a contest where participants were given gene expression datasets from four independent sources with the objective of predicting gene regulation. Analysing the results of DREAM5, the areas under precision recall curves (AUPR) were found to be around 10% for biological networks [18]. Such low performances pose a problem for interpretations of GRNs and ultimately hinder biological advances. Biological experiments are both time-consuming and costly, and performing experiments such as drug screening based on network predictions with an accuracy of 10% is ineffective.

However, there are some methods that have addressed the problem of low accuracy in predictions of gene regulation. One commonly applied approach to address the problem of correlated explanatory variables is to apply repeated subsampling of data followed by model training [24, 25], commonly referred to as bootstrapping. However, such an approach will not fully avoid identifying edges that cannot be rejected, and instead will include identifications across correlated explanatory variables [25]. Another notable approach to the problem of low prediction accuracy is the Robust Network Inference, RNI tool [26], which has been developed to omit identifications that cannot be rejected by any model [22]. However, RNI is not, as of the date of publication, a publicly available tool [22]. Furthermore, RNI has been shown to be much stringent, with few predictions being made when the signal-to-noise ratio (SNR) is low [24].

Herein, we present the LInear Profile LIKElihood (LiPLike), which is a novel algorithm to predict gene to gene regulation with high accuracy. LiPLike was conceptualised by merging the profile likelihood method for parameter confidence estimation [27] and GRN inference, but with two major differences. First, LiPLike assumes a regression model, which allows for substantially larger networks to be analysed. Second, LiPLike does not search for a confidence interval for a certain parameter, but instead only compares two key points of interest: the parameter value that minimises the residual sum of squares (RSS), and the point where this parameter equals zero. If there is an alternative linear combination of the explanatory variables that can model a dependent variable equally well, LiPLike will refrain from including such a parameter. In other words, LiPLike will only infer gene regulations with high confidence, and will avoid identifying edges that are not well-determined from data. In the case of several highly correlated explanatory variables that all can individually explain the expression data of a target, LiPLike makes no edge prediction, whereas the LASSO would typically include one at random [28] and the Elastic Net would include all [12].

We tested the sensitivity of LiPLike by applying data from the DREAM5 challenge and compared this with the 36 GRN inference methods analysed in the challenge. Instead of assessing the performance of a method based on the AUPR or receiver operating characteristic (ROC) curves, we evaluated the accuracy of top-ranked predicted edges for both LiPLike and the DREAM5 contestants. We found LiPLike to have higher accuracy than all DREAM5 methods in the biological networks, and better than average accuracy when benchmarking against the *in silico* generated data. Moreover, we found that LiPLike could successfully remove false positive identifications from GRN predictions of other methods, and recognised this feature to be useful whenever high accuracy GRN predictions are sought from gene expression data. Finally, to make LiPLike available to the community we built a Python toolbox available for download at https://gitlab.com/Gustafsson-lab/liplike.

## 2 Materials and Methods

### 2.1 Problem description

LiPLike is a novel GRN inference method to maximise the accuracy of gene regulatory predictions. LiPLike minimises the false positive prediction rate by not identifying potential regulators where a regulation can be replaced by a linear combination of one or several other explanatory variables (Fig. 1a). Specifically, the LiPLike algorithm was inspired by the profile likelihood method, used to estimate confidence intervals of estimated model parameters. Consider a system of independent variables *X* and dependent variables *Y*.

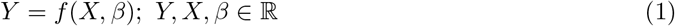

**Figure 1:**
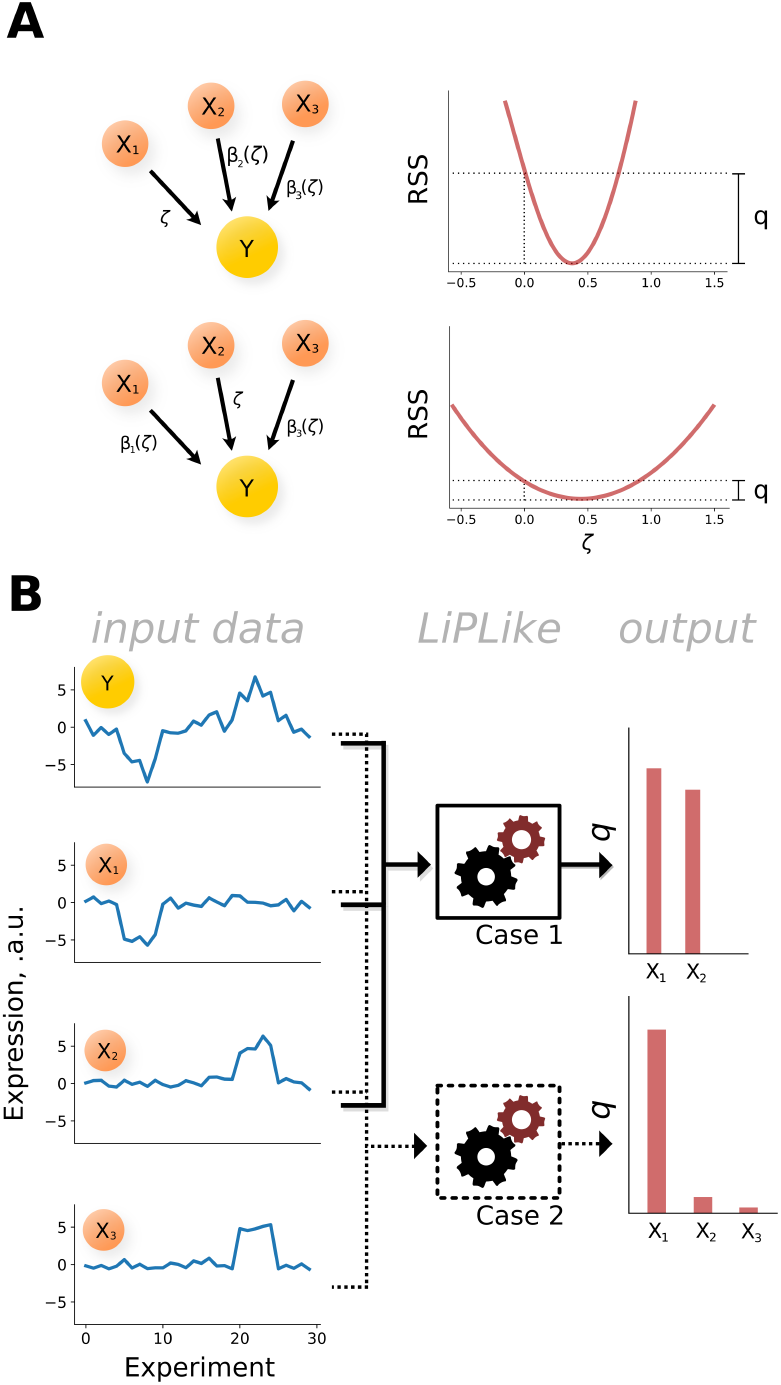
Illustration of LiPLike rationale. A) In a toy system of three gene regulators (*X*_1_, and the correlated variables *X*_2_ and *X*_3_) regulating a target gene (*Y*), the optimal parameters of the corresponding linear model are easily identified using the method of least squares. Next, by imposing a constraint that the parameter value of *X*_1_ should equal a value *ζ* and allowing for the remaining two to be adjusted accordingly, the profile of the RSS as a function of *ζ* can be studied. If *X*_1_ is uniquely needed to model *Y*, the RSS as a function of *ζ* will increase rapidly, as seen in the top case. In the bottom case there exists a linear combination of remaining explanatory variables, since *X*_2_ and *X*_3_ are correlated, and the RSS is less dependent on *ζ*. B) Two examples of LiPLike applied to data. Three independent variables exist, whereof two (*X*_2_ and *X*_3_) have a high correlation. To explain dependent variable Y, either (*X*_1_, *X*_2_) or (*X*_1_,*X*_3_) are needed, and there is no way to infer whether *X*_2_ or *X*_3_ is the correct regulator. If *X*_2_ or *X*_3_ is left out from the set, LiPLike infers all inputs to be important, as illustrated by the magnitudes of *q* shown to the right. When all three independent variables are included, LiPLike refrains from selecting variables that cannot be inferred uniquely.

In Equation (1), *β* is a vector of parameters mapping *X* to *Y*. The profile likelihood method estimates a confidence interval of a parameter *β*_*i*_ by observing the values of a variable *ζ*.

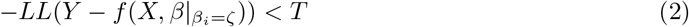

In Equation (2), the log-likelihood function [27], LL, is studied with respect to one parameter *β*_*i*_. By varying *ζ*, a confidence interval can be defined as the values of *ζ* for which the negative log-likelihood is below a threshold *T*. The profile likelihood is, however, mostly applied to non-linear systems and for grey-box models where all parameters are known to exist. Here, we instead assume that data are generated from a linear system of variables Y dependent on variables X via a vector of constants, *β*.

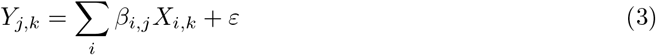

In the annotations of Equation (3), *Y*_*j,k*_ is a scalar corresponding to the expression of gene *j* at observation *k*. Likewise, *X*_*i,k*_ is a scalar corresponding to regulator *i* at observation *k*. As is widely known, for overdetermined systems the vector *β* can be analytically estimated to minimise the sum of squared residuals. Now, for each parameter *β*_*i,j*_, there are two parameter values that are of interest, namely the one minimising the RSS and the point where *β*_*i,j*_ = 0. Thus, we can quantify the relationship between these two points and introduce the term *q*_*i,j*_.

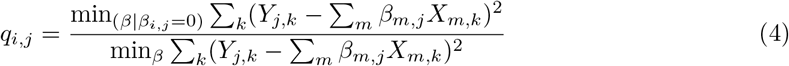

Given the values of Y and X, LiPLike returns *q*_*i,j*_ for all interactions in the network. As written in Equation (4), *q*_*i,j*_ will take larger values for edges that uniquely explain *Y*_*j*_ (Fig. 1b). In the case of two highly correlated regulators, *X*_*A*_ & *X*_*B*_, the constraint *β*_*a,j*_ = 0 will not independently give a considerable increase in the RSS. Thus, *q*_*a,j*_ ≈ 1, as *β*_*b,j*_ will adjust accordingly. In Equation (4), there are three properties that should be noted. First, if the system is not overdetermined, i.e. if rank(X) ≤ N, the denominator of Equation (4) equals zero, and thus *q* is undefined ∀*i, j*. To solve this problem, we implemented a functionality such that LiPLike either takes a prior interaction matrix as the input or builds one based on a cut-off on the Pearson correlation between variables. Second, when computing *q*_*i,j*_, the RSS of the special case of *β*_*i,j*_ = 0 is normed to the least squares fit of the fully connected system. Thus, in the case of a poor general fit to a dependent variable the increase in the numerator must thus be larger than for a well-fitted dependent variable for an edge to be identified. Third, the output of LiPLike *q*_*i,j*_, depends only on data properties and there are no additional hyperparameters that affect the performance of LiPLike.

### 2.2 Package design

We implemented the LiPLike algorithm as a Python toolbox based solely on built-in Python3 functions and the widely-used NumPy package. LiPLike generates ratios of the RSS, denoted *q*. To choose a cut-off for *q*, we implemented a Monte Carlo simulation of LiPLike applied on data randomly sampled from X and Y (default n=100 000). Next, we only considered LiPLike to have identified an edge where *q*_*i,j*_ takes a value outside the range of the Monte Carlo simulation. We also made this Monte Carlo simulation functionality part of the LiPLike package.

## 3 Results

### 3.1 LiPLike predicts edges with high accuracy

To test the ability of LiPLike to extract gene-gene interaction predictions, it was first applied to two versions of an *in silico*-generated gene expression dataset from the tool geneSPIDER [29]. The network contained 100 nodes genes with 300 measurements at SNR 10 and 100 000, respectively. We observed LiPLike to split edges into two groups, a group containing the majority of genes where no high precision prediction could be made (*q*_*i,j*_ ≈ 1), and a minor group where we had a clear indication of an interaction (*q*_*i,j*_ >> 1) (Fig. 2a).

**Figure 2:**
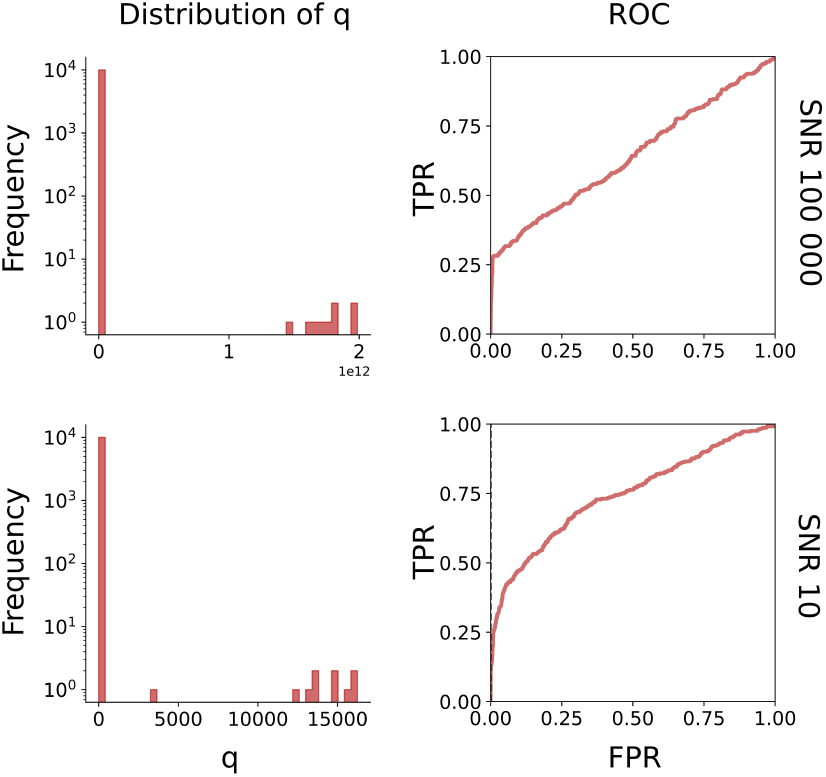
LiPLike properties and performance on in silico generated networks. The confidence of inferred edges is listed as q, calculated for two datasets for the same network. The networks differed in the signal-to-noise ratio (SNR). The magnitudes of q were found to be dependent on the SNR level, with a factor 10e7 differing between the datasets, as seen on the x-axis on the histograms to the left. Moreover, the histograms both display a property empirically arising from LiPLike for networks with strong signals, i.e. a separation of confidence into two distinct groups, high confidence or none. To the right are the corresponding ROC curves, showing that LiPLike infers edges well for some values, to then have a near to random chance for identifying an edge. The networks were retrieved from https://bitbucket.org/sonnhammergrni/genespider.

As expected we found that the spread between these two groups of edges appeared to be dependent on the SNR of the data, with an increase of *q*_*max*_ in order of 10^8^ when going from SNR=10 to SNR=100 000. Next, we studied the ROC curves for the respective datasets and observed a behaviour where the true precision rate was initially high, corresponding to a high accuracy. Next, the ROC curves transitioned into a linear phase, i.e. a phase where true predictions were randomly distributed. This linearity indicated that, as expected from Equation (4), LiPLike struggles to predict edges outside the high accuracy interval (Fig. 2b).

We further aimed at analysing the ability of LiPLike to handle under-determined systems, and sampled 50 experiments 100 times from the data with SNR=10, and applied LiPLike. By default, when applied to an under-determined network, LiPLike creates a prior network based on Pearson correlation between regulators and targets. In particular, for each gene LiPLike allows for n potential regulators, where n is 90% of the number of available observations. Using this approach, we observed a mean area under the ROC curve of 0.58, as compared to 0.75 when using the full dataset of 300 observations. This example showed how an over-determined system is not necessary for LiPLike to produce usable results, and that LiPLike can still be used when data are scarce.

### 3.2 LiPLike outperforms state-of-the-art methods in terms of accuracy

Knowing that LiPLike was capable of producing accurate predictions, we next aimed to benchmark LiPLike to other state-of-the-art GRN inference methods. There are several available datasets for benchmarking, whereof the DREAM5 network inference challenge is one of the most extensive. Moreover, the DREAM5 data contains ranked predictions of networks from both challenge participants, state-of-the-art methods, and a combined crowd estimate, all based on data from four different networks. The networks are based on two prokaryotic (*Staphylococcus aureus* & *Escherichia coli*), one eukaryotic (*Saccharomyces cereviciae*), and one *in silico* simulated system, all with accompanying expression datasets. However, due to the inability of DREAM5 contestants to infer edges from the *S. cereviciae* data [18], we excluded it from further analysis. We assumed the expression of target genes to be a linear combination of the transcription factors (TFs) in the data. Since the rationale of LiPLike is not to predict full networks, we measured performance in terms of accuracy, i.e. the rate of correct predictions among these top-ranked edges and the corresponding number of top-ranked edges in the DREAM5 contestants’ networks. Moreover, we identified the *E. coli* network to be of special importance, since the *S. aureus* gold standard was less extensive (only 518 edges), and since *in silico* is less relevant than true biologically derived networks.

Studying the performance of LiPLike on the *E. coli* data, we found accuracies of 0.38, 0.27, and 0.18 for 100, 500, and 1000 included edges respectively (Figure 3a-b). These numbers can be compared to the consensus model of all DREAM5 participants [18], with corresponding accuracies of 0.11, 0.10, and 0.08, i.e. less than half of LiPLike. Indeed, we found LiPLike to give the highest accuracy of all methods on the interval of 41-7943 of top-ranked edges. To choose a threshold of included edges, we performed a Monte Carlo re-sampling of data (Materials and Methods) and found 2308 gene regulations to have *q*_*i,j*_ larger than the max of the random *q* (*S. aureus*: 263, *in silico*: 2203). Studying the predictions of both DREAM5 biological networks we found LiPLike to have an accuracy better than all DREAM5 contestants at the Monte Carlo threshold, (LiPLike accuracy = 0.11, 0.10, for *S. aureus* & *E. coli*, which is an increase of 11% and 18% respectively compared to the best method in the DREAM5 challenge) (Fig 3c). Notably, applied to the biological datasets we found LiPLike to outperform the accuracy of the community predictions, which previously have been shown to be successful predictors of GRNs [18]. In the *in silico* generated data, we observed LiPLike to only generate a higher accuracy than average of the DREAM5 participants. It should be noted however that most methods performed well on this data, and the edges predicted by LiPLike still had an accuracy of 0.37 (average = 0.32). We further analysed the recall, i.e. the percentage of the edges in the gold standard that were correctly identified in the LiPLike top edges, and found the recalls to be 6%, 11%, and 32% for the *S. aureus*, *E. coli*, and *in silico* networks, respectively. With these results, we concluded that LiPLike, used as a stand-alone tool, is an effective method to extract gene-gene interactions with high confidence.

**Figure 3:**
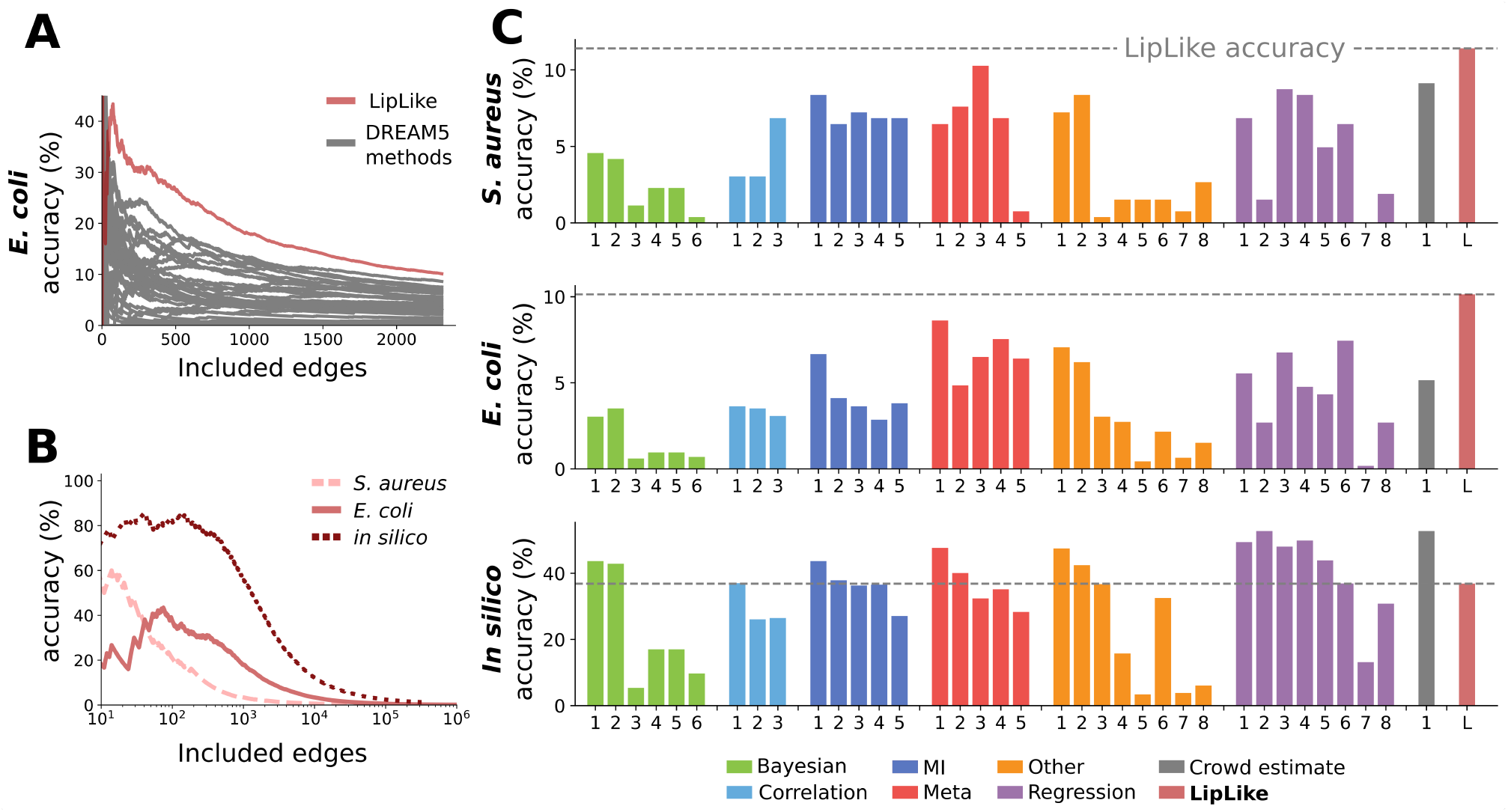
LiPLike performance on DREAM5 challenge data. A) Accuracy of algorithms predicting edges of the *E. coli* network as a function of number of edges considered. The LiPLike performance is plotted in red, showing a higher accuracy than all DREAM5 contestants. B) The accuracies of LiPLike across top-ranked edges for all networks. C) The accuracy of all methods, the crowd estimate, and LiPLike for the top-ranked edges. LiPLike gave the highest accuracy of all methods in both biologically derived networks, and ranked 20th of 36 in the in silico network.

### 3.3 LiPLike removed false positives from predictions of other methods

A major conclusion of the DREAM5 challenge was that merged method predictions robustly outperform individual methods. We therefore examined the impact of combining LiPLike with DREAM5 participants’ predictions. For the edges predicted by LiPLike and the same number of top-ranked edges in the community prediction (263, 2308, 2203 edges), we found significant overlaps (Odds ratio OR = 223, 288, 192, Fisher’s exact test *P* < 10^−74^, 10^−100^, 10^−100^ for the *S. aureus*, *E. coli*, and *in silico* networks, respectively). Indeed, we observed significant overlaps of edge predictions between LiPLike and almost all DREAM5 contestants (S1), with mean increases in accuracy being 161%, 105%, and 169% for the methods applied to the *S. aureus*, *E. coli*, and *in silico* data, respectively (S2).

Since the rationale of LiPLike is to not identify any edge where there is more than one potential regulator, we further tested to combine LiPLike with other methods for better accuracy and studied the DREAM5 community and LiPLike top-ranked predictions in depth. For the predictions in common for both methods, we found accuracies increased from 9%, 5%, and 53% to 53%, 15%, and 70%, (Fisher’s exact test *P <* 1.8 ⋅ 10^−14^, 5.4 ⋅ 10^−18^ 2.6 ⋅ 10^−78^ for *S. aureus*, *E. coli*, and *in silico* respectively (Fig. 4). Moreover, for the non-overlapping edges in the *E. coli* and *S. aureus* networks, we also observed an almost complete depletion of correct edges predicted by the community but not by LiPLike, with accuracies as low as 1.8% and 1.3% respectively (Fig 4), suggesting that combining LiPLike with other methods can effectively remove false positive edge identifications from a set of predictions. Furthermore, this property of removing false positive interactions seems to be a general property of LiPLike. By examining the intersects between all DREAM5 participants and LiPLike, we found the accuracy to be increased for almost all methods, with median increases in accuracy from 0.05, 0.04,and 0.36 to 0.47, 0.15, and 0.68, for *S. aureus*, *E. coli*, and *in silico*. Thus, there is a strong indication that LiPLike can be used in combination any GRN inference method to stratify predictions into more or less reliable interactions.

**Figure 4:**
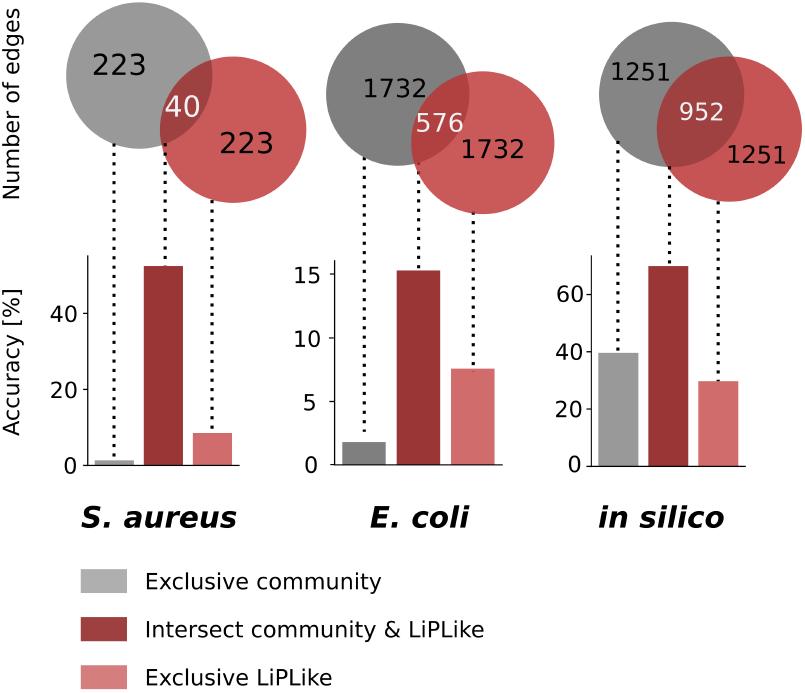
Accuracy of edge predictions of the DREAM5 community prediction and LiPLike, split up between top edges that are exclusively found in the community, LiPLike, and in both. In all cases the edges that are found in both predictions have a considerable increase in accuracy compared to the DREAM5 challenge community prediction. Moreover, in the case of the biological networks, *S. aureus* & *E. coli*, LiPLike performs better than the community in the non-overlapping predictions, indicating that LiPLike identifies edges that the community failed to include.

### 3.4 LiPLike robustly identifies interactions from a wide section of TFs

We next aimed to further compare the network structures generated by LiPLike and the community. Since LiPLike aims to only include edges that uniquely model an interaction, we first analysed the Pearson correlation of inferred regulators in the LiPLike and top community networks. For each inferred edge, we searched for the closest correlating TF and found the median of these correlations to be significantly higher for the community predictions than LiPLike for all tested datasets (median_*community*_ − median_*LiPLike*_ = 0.086, 0.106, 0.053, Mann-Whitney *P* < 10^−17^, 10^−124^, 10^−30^), for *S. aureus*, *E. coli*, and *in silico*, respectively (Fig 5a). Thus, LiPLike avoided identifying edges when two or more regulators correlated, which was not the case for the community predictions. It can be noted that the median Pearson correlation coefficient of the closest correlating TF in the three networks was significantly lower in the *in silico* generated network than in the experimentally generated ones (median_*S. aureus*_ = 0.61, median_*E. coli*_ = 0.54, median_*in silico*_ = 0.39, Mann-Whitney P_*in silico-S. aureus*_ < 10^−21^, *P*_*in silico-E. coli*_ < 10^−19^). It is possible that the lower Pearson correlations in the *in silico* data explains why LiPLike does not achieve a higher accuracy than the DREAM5 contestants, as it is thus rarer that two TFs can independently predict a gene expression.

**Figure 5:**
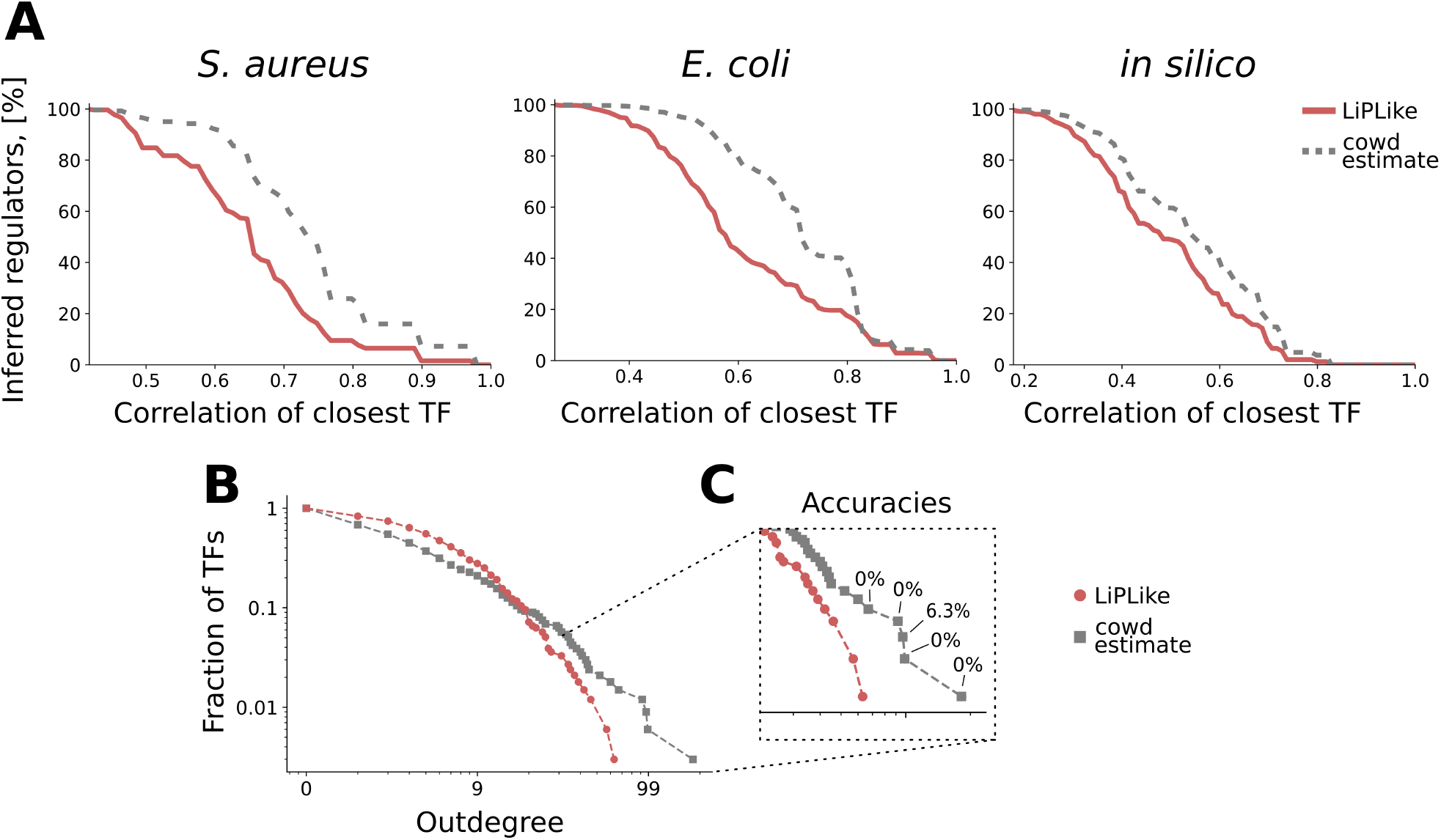
Cumulative distribution of the highest correlation with other regulators of putative interactions shown for LiPLike (red), and the crowd (grey) top-ranked interaction from respective DREAM5 networks. The regulators LiPLike identify tend to have on average fewer correlating regulators. For example, in the *E. coli* network we observed a median Pearson correlation of *ρ* = 0.57. For the corresponding community prediction, 85.3% of all inferred regulators have a higher correlation than 0.57 to another regulator. This higher correlation indicates that LiPLike to a lesser degree predicts edges where there are several potential regulators to choose from. B) Distribution of inferred edges for each transcription factor for LiPLike (red) and the community (grey). While the putative outdegrees of transcription factors in the community estimate appear to follow power law (as indicated by the straight line in the log-scale), LiPLike appears to select edges with a broader distribution profile. C) Accuracies for the inferred top regulators in the community prediction were found to be low. The top regulators in the LiPLike network had similar accuracies to the overall LiPLike accuracy.

Apart from predicting individual gene interactions, GRN inference can also be used to extract topological information on a biological system. For instance, network regulators with a disproportionately large outdegree have previously been used in predictive medicine [8]. We hypothesised that such genes are more likely to have closely correlated regulators, and that LiPLike therefore would fail to identify their targets. Studying the *E. coli* network, we found that LiPLike indeed predicted networks containing fewer high outdegree TFs than the community predictions. Whereas the community-predicted network followed power-law distribution on TF outdegrees, LiPLike produced a topology more like to an exponential distribution (Fig 5b). In other words, the DREAM5 community produced a network where, compared to LiPLike, the predicted edges to a greater extent congregated around a few transcription factors. We therefore tested whether these TFs were important regulators that LiPLike failed to capture. Surprisingly, we observed accuracies much lower than that of the overall community accuracy (0.01 compared to 0.05 overall accuracy), with four of the five top TFs having no predicted edges with support in the gold standard (Fig. 5c). As a comparison, the top five TFs predicted by LiPLike showed a combined accuracy of 0.09, in line with the 0.10 overall LiPLike accuracy, indicating that LiPLike can robustly predict edges across node outdegrees.

Lastly, we tested whether the top regulators in the different DREAM5 contestants overall had a low accuracy and continued to analyse all contestants’ regulators with a larger outdegree than any of the respective LiPLike regulators. Surprisingly, the *S. aureus* and *E. coli* networks on average had lower accuracies than the overall accuracy of the method (*S. aureus* and *E. coli*: 88 of 90 and 158 of 200 hubs having lower accuracies than the method overall performance, (binomial *P*_*S. aureus*_ < 10^−23^, *P*_*E. coli*_ < 10^−16^)), indicating that reduced accuracy in inferred edges from top gene regulators is a general property of GRN inference methods.

## 4 Discussion

The creation of gene regulatory networks is a pivotal tool for understanding gene expression patterns. However, such GRN inferences have long struggled with low accuracy in predictions, to a large extent due to correlating expression profiles of potential regulators. Herein, we have presented the LInear Profile LIKElihood (LiPLike), which is a novel method and software for high accuracy gene regulatory predictions from large biological datasets. As input, LiPLike takes data for dependent and independent variables and searches for cases where an independent variable uniquely explains the behaviour of a dependent variable. We showed that LiPLike could successfully infer gene-gene interactions from biological data, as a benchmarking towards the DREAM5 network inference challenge [18] and produced accuracies higher than all 36 DREAM5 contestants for GRNs of *S. aureus* and *E. coli*. We also reported that combining LiPLike with networks from other GRN prediction methods significantly increased the accuracy for gene-gene interaction predictions, indicating that LiPLike can be used to remove false positive predictions from GRN predictions of other methods.

LiPLike is related to the method profile likelihood, which aims to estimate the intervals that an estimated parameter can take with a retained fit to data [30]. However, LiPLike differs from the profile likelihood in three key aspects. First, the profile likelihood is most commonly applied to non-linear ordinary differential equation systems, while LiPLike assumes linear relations between independent and dependent variables, which increases the computation speed by several orders of magnitude. Second, the profile likelihood tests which values a model parameter, *β*, can take with retained fit to data. LiPLike only tests the parameter *β* for the values minimising the RSS and *β* = 0. Third, the profile likelihood estimates the uncertainty of a parameter, while the LiPLike orders all potential interactions in a network and interprets the increase in RSS from the two cases as an indication of present interaction.

While LiPLike is different from the profile likelihood in that it does not aim to infer edges in a network, there are alternative methods available for high accuracy GRN inference. The RNI method [26], for example, aims to only include edges that cannot be rejected by any model [22]. However, RNI might be too stringent and has been found not to make any interaction predictions from *in silico* generated data with SNR values commonly reported in biology [22, 31]. Another method aiming to address the problem of correlating explanatory variables is the random LASSO [25], which bootstraps the explanatory variables in a series of steps and predicts a network by taking the average results from the bootstrap outcome. Thus, correlated explanatory variables will be predicted a smaller number of times, but will still be predicted in the final outcome [25].

The rationale behind LiPLike is to only identify edges that cannot be replaced by other interactions in the data. This approach is different from other approaches, which often try to infer the most probable network. This difference makes LiPLike highly stringent, and it is therefore closer to be a method for edge identification than a tool for full network reverse engineering. Such identifications are important, for example, when planning costly follow-up experiments. Herein, we showed that LiPLike seems to have a higher accuracy than other available tools for edge identification when explanatory variables are highly correlated. We further hypothesised this performance to stem from the properties of common GRN inference methods. Indeed, when encountering correlated independent variables, GRN inference tools have been known to identify a regulator at random, or to include all potential regulators [12]. LiPLike, however, would identify none as a potential regulator. Importantly, we have also shown that LiPLike to be able to remove false predictions from GRN produced from other methods, a property that can be used by anyone that wishes to stratify their predictions into sets of high and low confidence.

## 5 Conclusion

Correlating explanatory variables poses a major obstacle when inferring gene regulatory networks. Available methods for GRN inference handle correlations by including several or one of the correlating variables. Herein, we presented LiPLike, which identifies no interactions that cannot be uniquely inferred from data, and we show LiPLike to predict edges with higher accuracy than other state-of-the art GRN inference tools in the DREAM5-challenge. Importantly, we also show that LiPLike can be used to remove false interactions from other methods, with the average increase of accuracies being 0.05, 0.04, and 0.36 to 0.47, 0.15, and 0.68, for *S. aureus*, *E. coli*, and *in silico*, respectively, and we recommend LiPLike be used on top of GRN estimations to give reliable predictions. In summary, we herein make gene interaction identification of high accuracy possible for the community, using LiPLike together with other algorithms or as a stand-alone feature selection tool.

## Acknowledgements

We would like to thank Andreas Tjärnberg for his input on *in silico* generated networks.

