## supplement figures for "LiPLike: Towards gene regulatory network predictions of high-certainty"

# S1

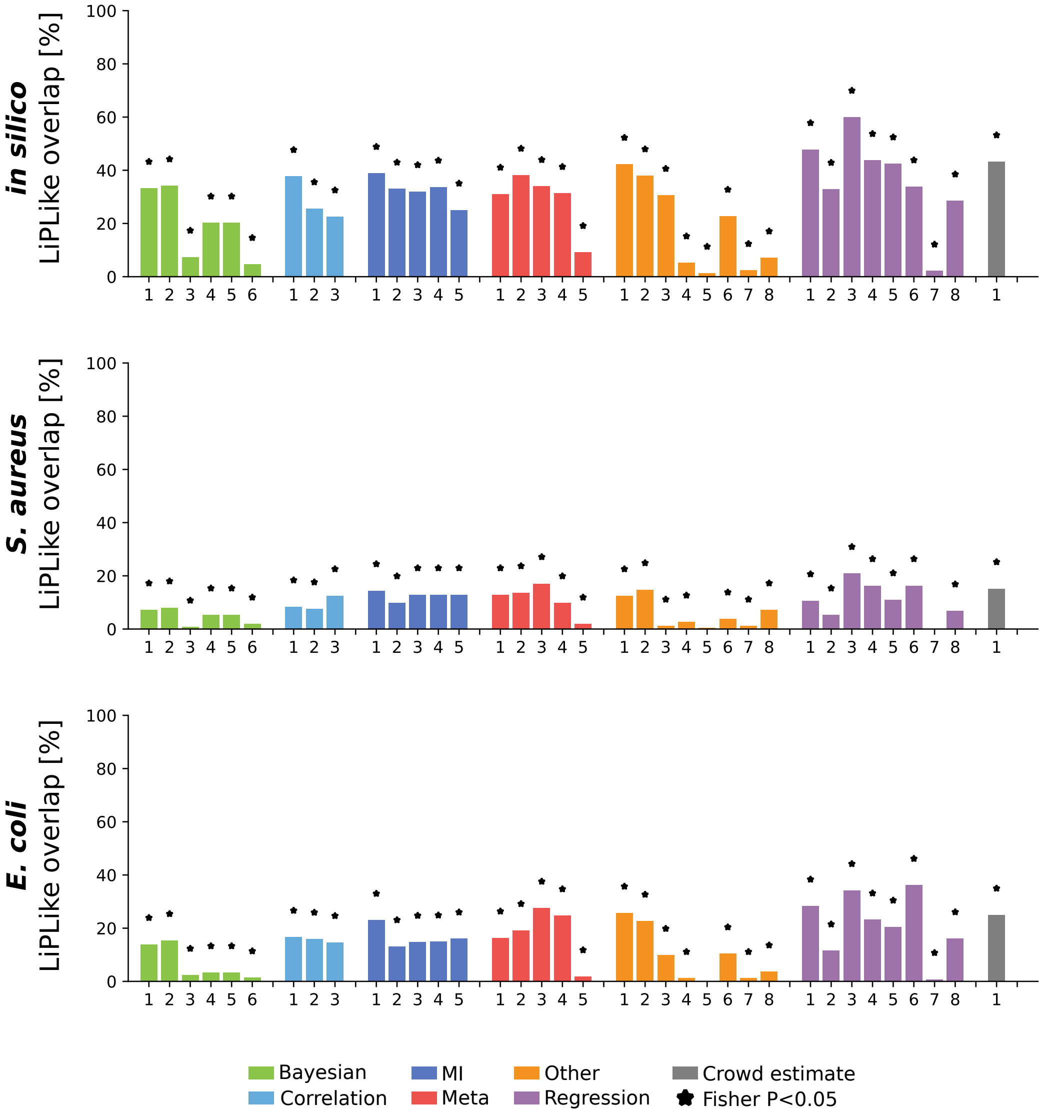


**S1:** Overlap between the DREAM5 method top edges and the corresponding LiPLike predictions. Almost all methods showed a significant overlap with LiPLike, with a general tendency for regression-based methods to identify edges in common.

# S2

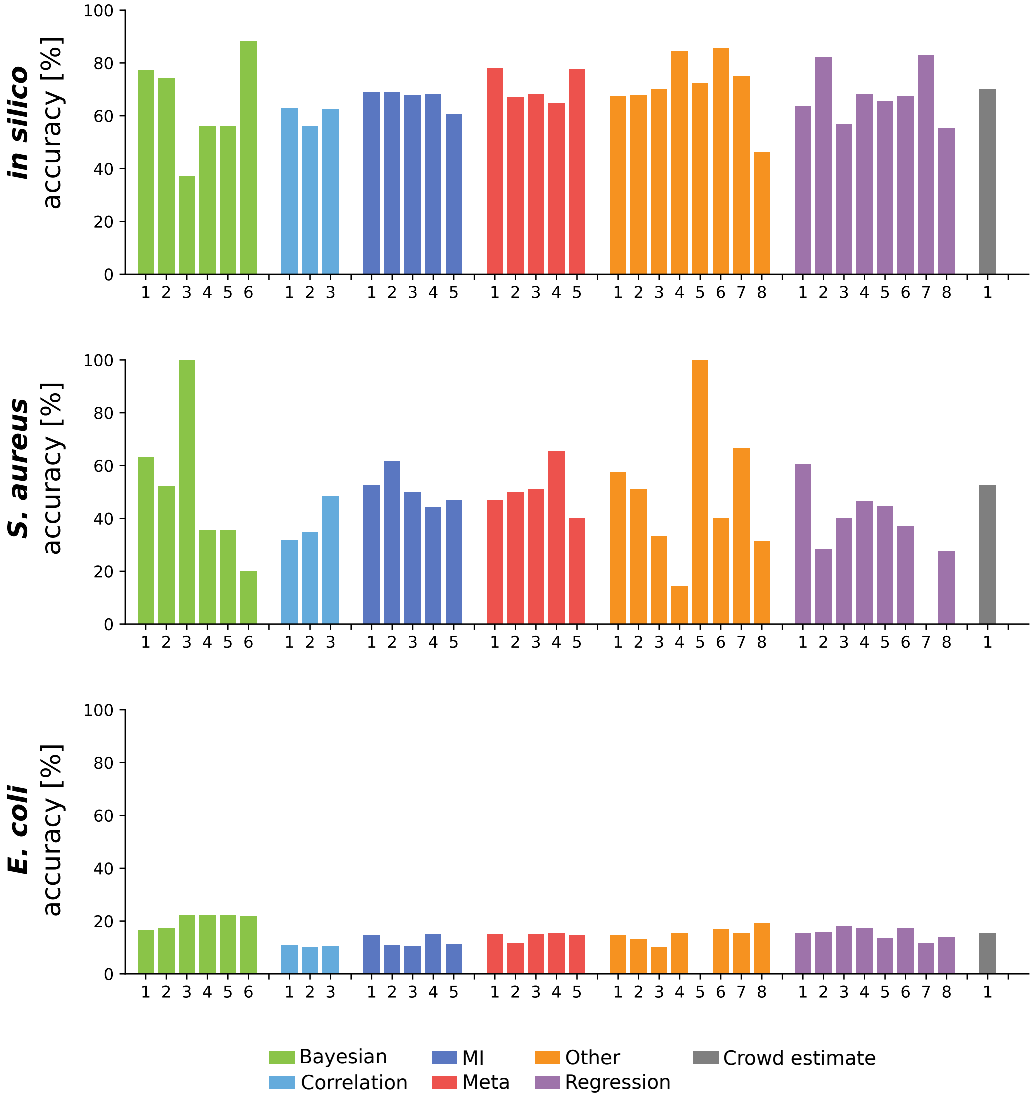


**S2**: Accuracies for the overlaps of top predictions between LiPLike and the DREAM5 challenge participants. Increases in accuracy were seen for almost all pairs. The number of prediction depends on the overlap, and thereby differs between each method and dataset.
